# Estimating invasion dynamics with geopolitical-unit level records: performance and similarity of common methods using both simulated data and a real case

**DOI:** 10.1101/2020.01.02.893248

**Authors:** Wanwan Liang, Liem Tran, Gregory Wiggins, Jerome Grant

**Affiliations:** Center for Geospatial Analytics, North Carolina State University, Raleigh, NC, USA; Department of Geography, University of Tennessee, Knoxville, TN, USA; National Institute for Mathematical and Biological Synthesis, Knoxville, TN, USA; Department of Entomology and Plant Pathology, University of Tennessee, Knoxville, TN, USA

**Keywords:** Invasion dynamics, spread pattern, geopolitical unit, county-level record, long distance jump dispersal, asymmetric spread

## Abstract

Estimating invasion dynamic is important to the management of invasive species, and geopolitical-unit level data are usually the most abundant and available records of invasive species. Here, for the first time we evaluated performances and similarities of eight common methods to estimate spread pattern and spread dynamic of invasive species with geopolitical-unit level data, and assessed impacts of variations in geopolitical-units on each method using simulated spread data. We also formulated a concave hull boundary displacement method (i.e., CEB) and an area-based regression method (i.e., AER) for estimating spread with geopolitical-unit data. Three regions with different sized counties in the United States (U.S.) were selected to conduct simulations and three spread scenarios were simulated. R^2^ and root mean square error were used to evaluate the abilities of all methods to estimate spread. Correlation coefficients were used to assess the similarity pattern of all methods. Finally, kudzu bug *Megacopta cribraria*, an invasive insect in the U.S., was used as a case study to test the generality of some results concluded from the simulated research. We found the CEB and two regression methods consistently estimated the right expansion patterns. Two boundary displacement and two area-based regression methods estimated highly correlated spread and were the best four methods, among which CEB had the best estimation. Distance-based regression methods are sensitive to irregularity and stochasticity in spread, and the minimum spread distance method had low ability to estimate spread. The case study showed consistent results with the simulated research. Both regression and boundary displacement methods can estimate spread patterns, overall rate, and spread dynamics of invasive species. Boundary displacement methods best estimate spread rates and dynamics; however, for spread without clear infestation outlines, area-based regression methods can be good alternatives.

## 1 Introduction

As a major component of global change, biological invasions have put growing threats on ecosystems and human society and caused economic loss globally (Paini et al., 2016, Pejchar & Mooney, 2009, Walsh et al., 2016). For example, the economic loss caused by invasive species was estimated at about $120 billion annually in the United States (U.S.) (Pimentel et al., 2005) and £1.7 billion in Great Britain (Williams et al., 2010). Gurevitch and Padilla (2004) summarized that among the species listed in the International Union for Conservation of Natural Resources Red List, 882 terrestrial species, 59 freshwater species, and 87 marine species are endangered by invasive species. Management of invasive species, therefore, becomes essential to minimize their negative impacts. Modeling invasion dynamic is important to the management of invasive species, as it facilitates prediction of spatial and temporal invasion risks, enhances early detection, and help to determine important factors affecting the invasions (e.g., Liang et al., 2019; Paini et al., 2016; Stohlgren & Schnase, 2006).

In practice, estimating invasion rate has been conducted on various species at all spatial scales including local (e.g., Haregeweyn et al., 2013, Sharov et al., 1999), regional (e.g., Evans & Gregoire, 2007), continental (e.g., Pyšek et al., 2008), and global scales (e.g. Suarez et al., 2001). Selection of invasion records to estimate spread is closely related to the spatial scales of research. For research conducted at local or smaller scales, spread data collected through field sampling or censuses are usually used (e.g., Pratt et al., 2003; Sharov et al., 1999). For plant species, time-series satellite or aerial images are also used (e.g., Haregeweyn et al., 2013; Pyšek et al., 2008). However, for research at larger scales, such as regional or continental scales, researchers usually have to collect all available records from online databases, published research, surveys, or field sampling (e.g., Masciocchi & Corley, 2013; Pyšek et al., 2008; Suarez et al., 2001). Consequently, data for large-scale research usually has coarse and non-unified resolution.

Collected data are often aggregated to a geopolitical-unit (i.e., city, county, parish, state, etc.), recording presence/absence of invasive species for each unit (e.g., Evans & Gregoire, 2007; Horvitz et al., 2017; Lantschner et al., 2014). Additionally, quarantines initiated by governments or institutions are usually conducted at the geopolitical-unit level, such as county or township (e.g., Perrins et al., 1993; Tobin et al., 2007 & 2015). Therefore, geopolitical-unit level data are usually the most abundant and available records of invasive species (Evans & Gregoire, 2007; Liebhold et al., 1992; Tobin et al., 2007; Tobin et al., 2015). During the past decades, researchers worldwide have used geopolitical-unit records to estimate invasion rates of various species (e.g., Evans & Gregoire, 2007; Horvitz et al., 2017; Perrins et al., 1993). Such research was mostly conducted at a regional scale (e.g., Horvitz et al., 2017; Morin et al., 2007; Perrins et al., 1993), as well as continental or global scales (e.g., Lantschner et al., Liu et al., 2014; 2014; Pyšek et al., 2008; Suarez et al., 2001).

Multiple methods have been commonly-used to estimate spread of invasive species over large spatial scales, and several researchers compared the accuracy of these methods to estimate spread of invasive species. However, to the best of our knowledge, this is the first research that evaluated performance and similarity of commonly used methods to estimate the overall invasion rates and invasion dynamics with geopolitical-unit level records using both simulated data and a case study. Existing research only compared the overall spread rate (Gilbert & Liebhold, 2010; Tobin et al., 2015), however, spread of invasive species, especially at large scales, is commonly complex due to spatial heterogeneity and stochastic events (Hastings et al., 2005; Pyšek et al., 2008). Estimating spreads with geopolitical-unit data further increases the uncertainties, as there can be large variations in the sizes of geopolitical units (Hastings et al., 2005; Pyšek et al., 2008). Consequently, compared to one single overall spread rate, estimating spread dynamics is more informative for understanding invasions (Hastings et al., 2005). Additionally, assessments of similarity patterns among common methods and their ability to estimate expansion patterns of invasive species are rare (e.g., Gilbert & Liebhold, 2010; Goldstein et al., 2019; Tisseuil et al., 2016; Tobin et al., 2015). Knowing the intrinsic similarity of commonly used methods on estimating spread patterns, i.e., the spatial or temporal spread dynamics, can further help to select method for calculating spread rates. Estimating expansion patterns of invasive species can facilitate prediction of further spread.

Currently, a systematic research on evaluating performances and similarities of common methods on estimating both overall spread rate and spread dynamics with geopolitical-unit level records, while considering variation and stochasticity in real landscape is lacking. Thus, this research aimed to address these gaps. Specifically, we conducted evaluations from the following perspectives: 1) the accuracy of commonly-used methods to estimate expansion pattern, overall rate, and spread dynamics with geopolitical-unit level records while considering anisotropy and stochasticity in spread, 2) the impact of the size of geopolitical-units on each method, and 3) the intrinsic similarity pattern of common methods to estimate spread. We used simulated data, as spread rate and dynamics can be accurately calculated from simulations. We also formulated an alternative boundary displacement method and area-based regression method for estimating spread with geopolitical-unit data. To test the generality of results from the simulated research on the similarity pattern among common methods, we used an invasive species in the U.S., kudzu bug, *Megacopta cribraria* (Heteroptera: Plataspidae), as a case study. It’s worth noting that this research aimed to evaluate commonly-used methods to estimate spread with geopolitical-unit level records. Nevertheless, alternative approaches are also available with geopolitical-unite level record (e.g., Fitzpatrick et al., 2010; Goldstein et al., 2019; Meentemeyer et al., 2011; Pioz et al., 2011).

## 2 Common Methods to Estimate Spread with Geopolitical Unit

### 2.1 Regression Methods

The most commonly-used methods to estimate spread are regression and boundary displacement methods, which can be used with all types of invasion records (summarized in Gilbert & Liebhold, 2010; Tobin et al., 2015). The general idea of regression methods is to regress the spread measurement against time, beginning when the infestation is first observed.

#### Spread Distance

With geopolitical-unit records, one way to calculate spread distance is to derive the minimum distance between the spread origin and the polygon of each geopolitical unit (Tobin et al., 2007 & 2015). Distances between the spread origin and centroids of infested geopolitical units also have been used (Evans & Gregoire, 2007; Liang et al., 2019).

#### Square Root Area

This method assumes an invasive species is spreading by approximately concentric circles, for which the total spread distance (*D*) can be estimated as 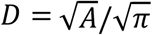, where A represents the cumulative area of the spread regions (Shigesada & Kawasaki, 1997; Skellam, 1951). For geopolitical-unit level records, the square root of the cumulative area of all infested geopolitical units is used as the measurement (Tobin et al., 2015).

#### Number of Infested Geopolitical Unit

Directly regressing the cumulative number of infested geopolitical units, *n*, by invasion times has been used in previous research (e.g., Perrins et al., 1993; Pyšek et al., 2008; Suarez et al., 2001). However, we argue that the square root of the cumulative number of infested units, 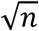, should be used instead (Williamson et al., 2005). If the total number of infested geopolitical units is *n* and the mean county size is Ā, then, the total infested area is *A = n*Ā. As mentioned above, the spread distance can be estimated as 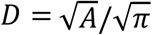, and by replacing *A* with *n*Ā we get 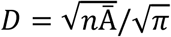. Therefore, *D* is linearly associated with 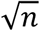. The spread rates estimated from 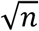 should be linearly correlated with the ones by spread distance regression and square root area regression. Additionally, the measurement 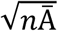 can be an alternative to the 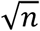 method and be used as an area-based regression method.

### 2.2 Boundary Displacement Method and Minimum Spread Distance Method

Boundary displacement methods estimate spread rates as the distance between consecutive infestation limits from different time periods. The outer boundary of geopolitical units infested within the same period is commonly-used as the infestation outline (e.g., Liebhold et al., 1992; Tobin et al., 2015). This method may require slight changes to the infestation boundaries to avoid folds, islands, and gaps on the boundary (Sharov et al., 1999). We also used the concave hull method to derive the infestation boundaries by using polylines that connect the centroids of all the outermost newly-infested geopolitical units as the boundary. This method avoids the need to modify infestation boundaries, and the infestation boundaries can be derived automatically using programs such as R (see Appendix S1).

MSD takes the minimum distance between a newly-infested geopolitical unit and all units infested in earlier periods as the distance that a species has to spread to invade the new unit (Aikio et al., 2010; Horvitz et al., 2017). The mean of all minimum distances of geopolitical units infested in the same period is taken as the spread rate in that period. Similar to boundary displacement methods, MSD also directly estimates temporal spread dynamics.

## 3 Methods

### 3.1 Spatial Area of Simulated Spread

To represent the real-world variety of geopolitical units for the generality of the simulated research, we used the counties in the U.S. for the simulation study (Fig. 1 (a)). The county size distribution in the U.S. is representative of many countries where the size of geopolitical units varies largely among different regions (e.g., Canada, China, Mexico). To evaluate the performance of common methods to estimate spread with different geopolitical-unit sizes, we conducted simulations individually in three regions, Region 1 (R1), Region 2 (R2), and Region 3 (R3) (Fig. 1 (a)). In each region, we set a spread origin, from which the spread was simulated. The county size and its coefficient of variation (CV) in three regions are listed in Table 1.

**Fig. 1.**
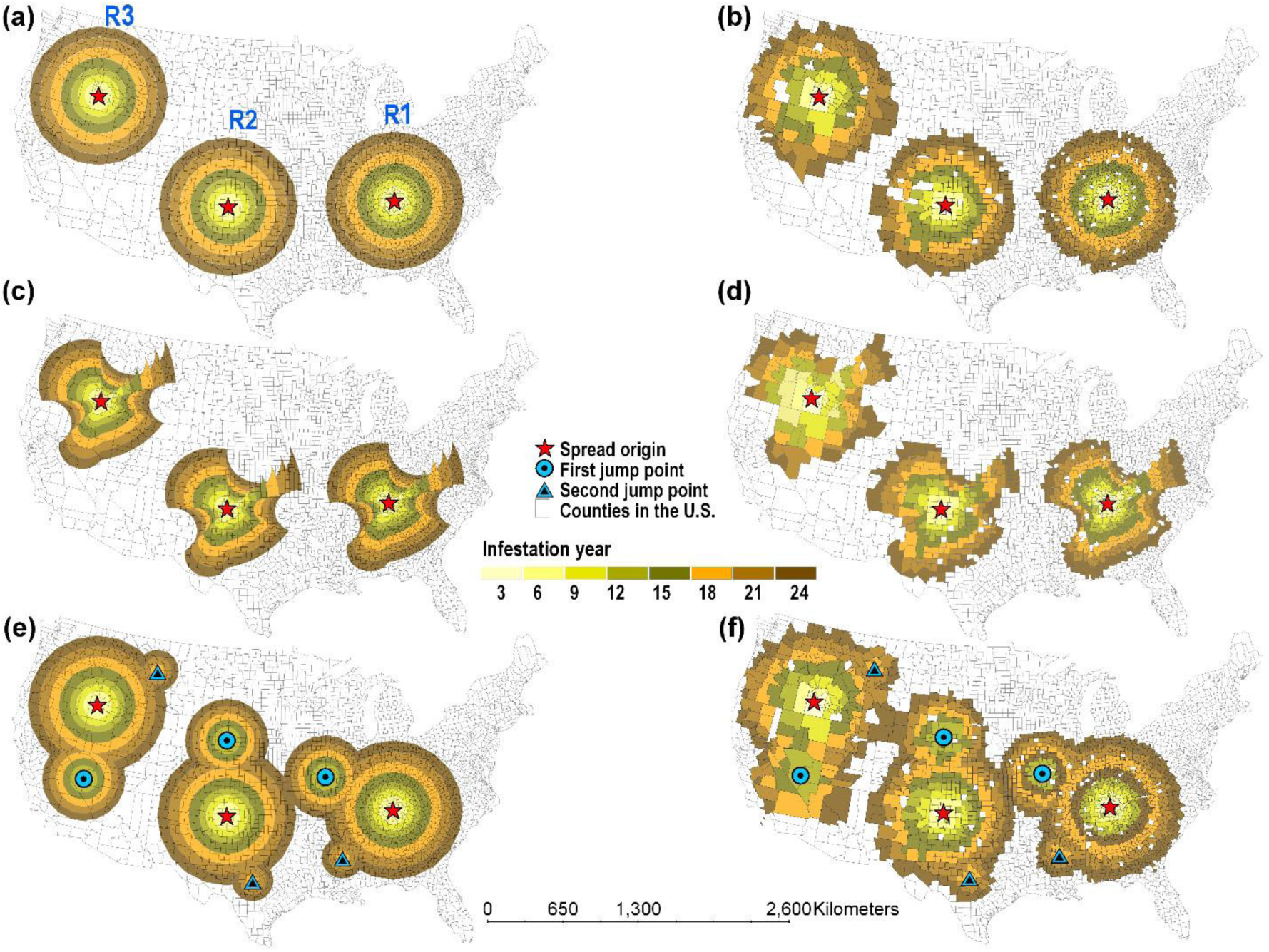
Example of simulated invasion dynamics and conversion to county-level data using Type 2 spread pattern (biphasic spread). (a), (c), and (e) Simulated biphasic spread during years 3-24 for scenarios 1, 2, and 3, respectively; (b), (d), and (f) county-level spread record converted from (a), (c), and (e), respectively. The distances between the spread origin and first and second jump points in (e) and (f) are set the same in all three regions.

**Table 1.** Statistics of county size in each region for different scenarios and types of spread.

| Spread scenarios | Spread type | No. of infested counties |  |  | Mean county area (km<sup>2</sup>) |  |  | Coefficient of variation |  |  |
| --- | --- | --- | --- | --- | --- | --- | --- | --- | --- | --- |
|  |  | R1 | R2 | R3 | R1 | R2 | R3 | R1 | R2 | R3 |
| S1 | Type 1 | 637 | 240 | 103 | 1188.21 | 3173.15 | 7564.29 | 0.12 | 0.07 | 0.29 |
|  | Type 2 | 919 | 371 | 172 | 1235.23 | 3133.17 | 7243.13 | 0.15 | 0.07 | 0.31 |
|  | Type 3 | 930 | 377 | 172 | 1236.82 | 3175.75 | 7229.18 | 0.16 | 0.10 | 0.32 |
| S2 | Type 1 | 516 | 198 | 82 | 1125.04 | 3133.54 | 8472.74 | 0.02 | 0.09 | 0.29 |
|  | Type 2 | 760 | 209 | 95 | 1176.03 | 3231.86 | 8062.71 | 0.06 | 0.07 | 0.33 |
|  | Type 3 | 758 | 209 | 95 | 1175.65 | 3231.86 | 8062.71 | 0.07 | 0.13 | 0.26 |
| S3 | Type 1 | 914 | 381 | 146 | 1271.61 | 3273.84 | 8858.33 | 0.12 | 0.04 | 0.35 |
|  | Type 2 | 1147 | 471 | 200 | 1290.96 | 3275.94 | 8169.05 | 0.13 | 0.04 | 0.35 |
|  | Type 3 | 1158 | 479 | 200 | 1288.32 | 3274.22 | 7881.00 | 0.14 | 0.08 | 0.34 |

S1, S2, S3, and No. are short for symmetric spread, asymmetric spread, long distance jump dispersal, and number, respectively.

### 3.2 Simulation of Three Spread Patterns and Three Spread Scenarios

To evaluate abilities of common methods to estimate spread patterns of invasive species, we simulated three types of spread summarized by Shigesada et al. (1995): 1) linear spread, 2) biphasic spread resulting from two linear-spread phases, and 3) logistic growth function spread. These three types of spread were commonly observed in research focused on invading organisms (e.g., Lantschner et al., 2014; Mineur et al., 2010). We simulated three spread scenarios to compare the performance of all methods. We first simulated a symmetric spread (S1) to evaluate accuracies of all methods under this ideal scenario (Fig. 1 (a)-(b)). To evaluate capability of all methods to deal with anisotropy in spread, we simulated an asymmetric spread (S2) reflecting heterogeneity (Fig. 1 (c)-(d)). We also simulated a LDJD (S3) to see how different methods response to this random event (Fig. 1 (e)-(f)). However, estimating the spread ability of invasive species with LDJD is arguable, as LDJD, which is caused by rare random event, can dramatically falsely increase the spread rate. Thus, we only simulated a scenario of LDJD, and more (probably better) methods to estimate spread rates with LDJD are included in Discussion. We simulated these three spread patterns for all scenarios and regions.

#### Simulation of Symmetric Spread

For the simulation of S1, the Type 1 has a constant rate for all periods. We set this rate to 20 km/year, as it approximates the mean spread rate of invasive species based on multiple research (e.g., Horvitz et al., 2017; Suarez et al., 2001; Tobin et al., 2007). For Type 2 the simulated rate was set to 20 km/year for the first 12 invasion years and 30 km/year for the following 12 years, thus the mean spread rate is 25 km/year for the whole period. The Type 3 follows a logistic growth function *y* = 826/(1 + *e*^-0.43∗(*x*-10)^), where *y* and *x* represent the total spread distance and spread time, respectively.

#### Simulation of Asymmetric Spread

Simulation of the three spread types for S2 is similar with S1, except that the rates varied among different directions (Fig. 1 (c)-(d)). The mean simulated rate in all directions varied between 10-24 km/year for Type 1, and 12-31 km/year for Type 2 and Type 3 spreads.

#### Simulation of Long-Distance Jump Dispersal

To simulate S3, we added two jump dispersal events in S1 with one occurring in year 9 and another occurring in year 18 (Fig. 1 (e)-(f)). To make the S3 in three regions comparable, the distances among the two jump points and the spread origin are set the same for all regions (Fig. 1 (e)-(f)). The jump point would become a new spread origin, from which further spreads occur in all directions. We set this rate to 20 km/year for all jump points and spread types for clarity and simplicity.

#### Simulated Rates

The Equidistant conic map projection was used for S1 and S3 to maintain the same spread distance in all regions, whereas the Lambert conformal conic map projection was used for S2 to maintain the asymmetry of spread in all regions. Unlike S1 (Fig. 1 (a)), the spread rates for S2 and S3 cannot be directly derived from the spread algorithm. The spread rates for S2 and S3 were derived as the mean spread distance in all directions in the given period (Fig. 1. (c), (e)). We also used this method to estimate the simulated rate for S1 to see how well the calculated rates match the simulated rates.

#### Converting Simulation to Geopolitical-Unit Level Spread

To convert the simulated spread (shown in Fig. 1 (a), (c), (e)) to geopolitical-unit level spread (shown in Fig.1 (b), (d), (f)), we selected counties that were infested in the same periods (i.e., every three years). A county is only defined as first infested when more than 10% area is included in the simulated spread zone to eliminate margin effect. To reflect real world stochasticity, we randomly assigned 5% of the counties first infested in each period as non-infested counties (Fig. 1).

### 3.3 Estimating Overall Rate and Spread Dynamics

Five regression methods, two boundary displacement methods, and a MSD were included in this research (Table 2). For boundary displacement methods, distances between two consecutive boundaries were measured as the mean length of all transects radiating from the spread origin at 2° (Liebhold et al., 1992; Liang et al., 2019). Technical flow of the simulation is shown in Fig. 2.

**Table 2.**
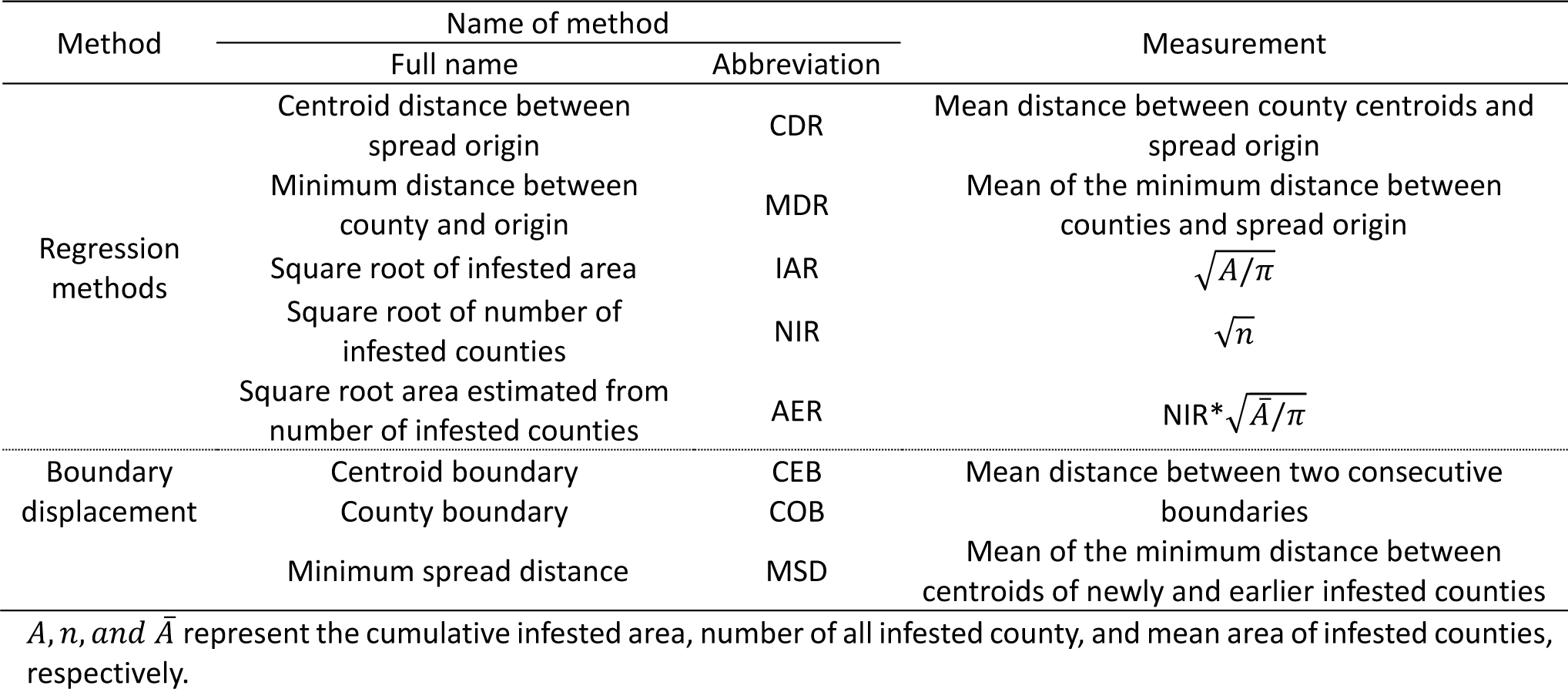
Full and abbreviated names of all methods and the measurements used by all methods to estimate spread dynamics

**Fig. 2.**
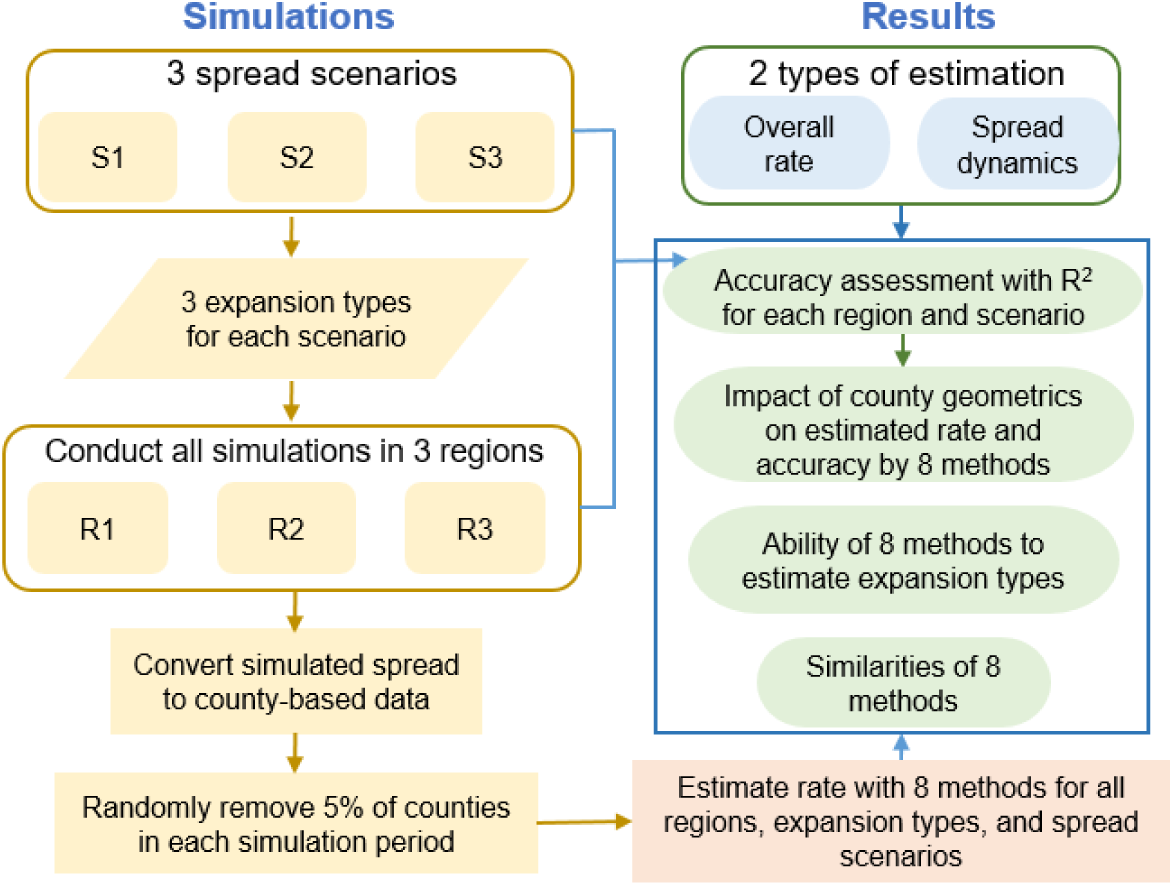
Technique flow diagram of the simulation study. S1, S2, and S3 represent symmetric spread, asymmetric spread, and long distance jump dispersal, respectively.

CEB, COB, and MSD directly estimate spread dynamics, and the overall rate was calculated as the mean of all periods. Spread dynamics were estimated as the difference of measurements between two consecutive periods for regression methods, and the overall rate for regression methods were estimated as the slope of a linear model for Type 1 and mean of two slopes of a segmented linear model with break point at year 12 for Type 2. For Type 3, instead of estimating spread rate using derivatives of the logistic growth function, we used the below model to make the estimation was more representative of the practice:

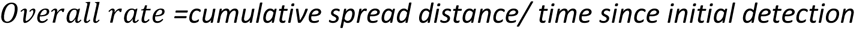

### 3.4 Evaluation Statistics

#### 3.4.1 Ability of All Methods to Estimate Spread Patterns

For CEB, COB, and MSD, we derived the cumulative values of spread measure for each period. To test whether all methods can accurately estimate the spread patterns, we fitted three regression models, i.e., linear, biphasic linear, and non-linear with logistic function, to the estimated spread measure for each spread type, region, and scenario. The Akaike information criterion (AIC) of three regression models were derived and the model with lowest AIC was assigned as the estimated spread pattern by each method (Akaike, 1973). For Type 1, we defined the method can accurately estimate the spread pattern if the linear or biphasic linear fit has the lowest AIC, as biphasic linear can be an overfitting of linear model but have lower AIC.

#### 3.4.2 Accuracy and Similarity of All Methods

We used paired t-test to assess the significance of difference between simulated and estimated overall rate by each method. We also derived R^2^ and the root mean square error (RMSE) between the estimated and simulated spread to evaluate the performance of each method on estimating both overall rate and spread dynamics. R^2^, varying from 0 to 1, is scale-independent with value 1 indicating perfect estimation of the simulated spread dynamics. RMSE measures the absolute deviation of estimation from the simulation, thus cannot be used on NIR method due to its sensitivity to the absolute value of estimated spread rates.

We derived the Pearson correlation coefficient (r) of derived spread rates among all methods to assess their similarity pattern on estimating spread. Hierarchical clustering (HC) with complete linkage was used to group all methods based on their similarity of derived spread rates.

#### 3.4.3 Impact of County Size on Estimation Accuracy

To determine whether the county size and its variation affect the values of estimated rates by each method, we tested the significance of correlations between the estimated rates with the mean and coefficient of variation (CV) of county size. To assess the impact of county size and its variation on accuracies of estimated rates, we tested the significance of correlations between the mean and CV of county size and R^2^ of the estimation for each region and spread scenario.

### 3.5 Case Study on Kudzu Bug

Kudzu bug had a phenomenal spread in the U.S., and has been reported in 652 counties by the end of 2017 (Eger et al., 2010; Liang et al., 2019). As a case study without accurate sampling data, the actual spread rate of kudzu bug is unknown. Therefore, kudzu bug was selected as a case study to compare the derived similarity pattern of common methods with the pattern from the simulated research. All eight methods were used to estimate the overall spread rate and spread dynamics of kudzu bug during 2010-2016. As kudzu bug had an asymmetric spread, we divided the infested area into eight neighborhoods and derived spread rates in all neighborhoods to better estimate its spread dynamics. Details on neighborhood classification of infested regions by kudzu bug can be found in Liang et al. (2019). The logistic growth function (Type 3) was used to estimate the rates for regression methods, as former research suggested Type 3 better delineated the spread pattern of kudzu bug (Liang et al., 2019).

## 4 Results

### 4.1 Ability of All Methods to Estimate Spread Patterns

Both regression methods and boundary displacement methods can estimate the spread patterns (Appendix S2). CEB, NIR, and AER correctly estimated the spread patterns for all scenarios and regions (see Appendix S2 for AICs). For S1, except MSD, all other methods correctly estimated the spread patterns (see Appendix S2). For S2 and S3, IAR and COB correctly estimated all spread patterns for R1 and R2, whereas MDR and MSD constantly misclassified the spread patterns for S2 and S3 (see Appendix S1).

### 4.2 Accuracy of All Methods

#### 4.2.1 Accuracy to Estimate Overall Spread Rate

The mean spread distance in all directions can provided an accurate measurement on the simulated rates according to its estimation for S1 (R^2^=1.00). For all scenarios, the MSD method consistently estimated a higher spread rate in regions with larger county size leading to underestimation of rates in R1 but overestimation in R3 (Table 3). For S1, all other methods estimated similar spread rates with simulated rates (Table 3). However, the MDR method estimated a significantly lower rate (*P*=0.003) and the COB estimated higher rate than the simulations (*P*=0.002). For S2 and S3, all regression methods tended to estimate significantly higher rates than the simulations (Table 3), whereas the boundary displacement methods estimated significantly higher spread rates when LDJD occurred.

**Table 3.** Simulated and estimated overall spread rates and p-values for the paired t-test between simulated and estimated rates for each spread scenario

| Spread scenario | Type | Simulated rate (km/y) | Region | Estimated Rates |  |  |  |  |  |  |  |
| --- | --- | --- | --- | --- | --- | --- | --- | --- | --- | --- | --- |
|  |  |  |  | CDR | IAR | MDR | NIR | AER | COB | CEB | MSD |
| S1 | I | 20.00 | R1 | 19.81 | 19.45 | 19.53 | 1.02 | 20.23 | 20.13 | 20.11 | 15.91 |
|  |  |  | R2 | 19.86 | 19.91 | 19.61 | 0.61 | 19.26 | 19.93 | 20.55 | 20.54 |
|  |  |  | R3 | 20.20 | 19.80 | 19.82 | 0.38 | 18.25 | 20.12 | 21.57 | 27.99 |
|  | II | 25.00 | R1 | 24.90 | 24.34 | 24.38 | 1.16 | 22.56 | 25.21 | 25.87 | 18.46 |
|  |  |  | R2 | 24.90 | 24.71 | 24.48 | 0.78 | 24.79 | 24.97 | 25.43 | 22.55 |
|  |  |  | R3 | 24.95 | 24.25 | 24.60 | 0.52 | 25.52 | 25.16 | 26.28 | 29.07 |
|  | III | 25.20 | R1 | 25.15 | 25.17 | 24.01 | 1.27 | 25.20 | 25.06 | 25.23 | 19.07 |
|  |  |  | R2 | 25.24 | 25.70 | 23.70 | 0.81 | 25.75 | 25.12 | 25.73 | 23.15 |
|  |  |  | R3 | 25.60 | 26.42 | 23.48 | 0.55 | 26.38 | 25.38 | 26.56 | 28.78 |
|  |  | *P* |  | *0.987* | *0.661* | *0.003* | *NA* | *0.468* | *0.260* | *0.002* | *0.742* |
| S2 | I | 16.68 | R1 | 18.00 | 16.89 | 17.84 | 0.88 | 16.70 | 16.42 | 16.13 | 15.15 |
|  |  | 16.59 | R2 | 19.03 | 17.41 | 18.82 | 0.54 | 17.01 | 16.85 | 17.18 | 20.19 |
|  |  | 16.59 | R3 | 19.72 | 17.75 | 19.09 | 0.34 | 17.73 | 17.16 | 17.42 | 29.09 |
|  | II | 21.44 | R1 | 23.54 | 21.14 | 23.09 | 1.06 | 20.49 | 21.08 | 20.81 | 17.56 |
|  |  | 21.33 | R2 | 24.47 | 22.61 | 24.41 | 0.70 | 22.50 | 21.90 | 22.23 | 22.21 |
|  |  | 21.32 | R3 | 23.95 | 21.37 | 23.57 | 0.45 | 22.64 | 20.36 | 21.14 | 29.34 |
|  | III | 21.42 | R1 | 24.04 | 21.86 | 23.08 | 1.13 | 21.83 | 21.04 | 20.65 | 17.88 |
|  |  | 21.31 | R2 | 24.63 | 22.92 | 23.11 | 0.73 | 23.50 | 22.05 | 21.87 | 22.82 |
|  |  | 21.30 | R3 | 25.97 | 23.30 | 24.09 | 0.48 | 24.54 | 21.67 | 21.17 | 28.22 |
|  |  | *P* |  | *0.000* | *0.005* | *0.000* | *NA* | *0.016* | *0.748* | *0.744* | *0.175* |
| S3 | I | 24.66 | R1 | 26.31 | 25.99 | 25.94 | 1.28 | 25.72 | 25.66 | 25.45 | 20.79 |
|  |  |  | R2 | 26.13 | 26.71 | 25.90 | 0.81 | 26.16 | 26.57 | 26.84 | 26.03 |
|  |  |  | R3 | 25.13 | 26.53 | 24.70 | 0.48 | 25.44 | 25.40 | 25.79 | 33.51 |
|  | II | 28.07 | R1 | 31.32 | 28.32 | 30.73 | 1.33 | 26.88 | 29.37 | 30.05 | 22.32 |
|  |  |  | R2 | 32.76 | 29.06 | 32.46 | 0.89 | 28.77 | 29.71 | 30.32 | 26.96 |
|  |  |  | R3 | 29.12 | 29.80 | 28.32 | 0.58 | 29.42 | 28.70 | 30.18 | 33.61 |
|  | III | 28.21 | R1 | 29.70 | 28.71 | 28.53 | 1.42 | 28.71 | 27.88 | 28.51 | 21.76 |
|  |  |  | R2 | 30.04 | 29.44 | 28.48 | 0.91 | 29.44 | 28.77 | 30.19 | 25.76 |
|  |  |  | R3 | 28.36 | 29.51 | 25.98 | 0.59 | 29.51 | 28.87 | 30.54 | 31.56 |
|  |  | *P* |  | *0.003* | *0.000* | *0.087* | *NA* | *0.009* | *0.002* | *0.000* | *0.976* |

S1, S2, and S3 represent symmetric spread, asymmetric spread, and long-distance jump dispersal, respectively. *P*: *P* value of paired t-test between estimated and simulated rates by each scenario (one-tale test for $P < 0.10$ , two-tale test for $P \geq 0.10$ ).

Based on R^2^, all methods estimated S1 best and S3 least (Table 4). The MSD and NIR methods estimated spread poorly when all three regions were analyzed (R^2^ <0.1). For S1 and S2, the CEB, CDR and MDR were the best three methods, whereas COB, CEB, and IAR were the best three methods for S3. Despite the large county sizes and great variations in the size of the counties in R3, all methods had higher R^2^ than that for S3. Based on both R^2^ and RMSE on spread rate for all scenarios and regions, CEB had the best estimation, followed by COB, IAR, and AER.

**Table 4.** R^2^ of estimated overall spread rate by each scenario and region, and R^2^ and root mean square error (RMSE) of estimated spread rates in all regions for all scenarios

| Spread |  | Regression Method |  |  |  |  | Boundary Displacement |  | MSD |
| --- | --- | --- | --- | --- | --- | --- | --- | --- | --- |
|  |  | CDR | IAR | MDR | NIR | AER | CEB | COB |  |
| $R^2$ by scenario | S1 | 0.99 | 0.95 | 0.96 | 0.09 | 0.87 | 1.00 | 0.95 | 0.04 |
|  | S2 | 0.93 | 0.92 | 0.96 | 0.09 | 0.86 | 0.94 | 0.93 | 0.03 |
|  | S3 | 0.72 | 0.89 | 0.47 | 0.02 | 0.79 | 0.85 | 0.89 | 0.00 |
| $R^2$ by region | R1 | 0.91 | 0.98 | 0.89 | 0.93 | 0.92 | 0.98 | 0.98 | 0.89 |
|  | R2 | 0.84 | 0.96 | 0.80 | 0.96 | 0.95 | 0.97 | 0.98 | 0.76 |
|  | R3 | 0.82 | 0.95 | 0.81 | 0.92 | 0.91 | 0.98 | 0.97 | 0.41 |
| All rates | $R^2$ | 0.85 | 0.95 | 0.81 | 0.14 | 0.91 | 0.98 | 0.97 | 0.13 |
|  | RMSE | 2.13 | 1.06 | 1.78 | NA | 1.25 | 0.71 | 1.22 | 5.19 |

S1, S2, and S3 represent symmetric spread, asymmetric spread, and long-distance jump dispersal, respectively.

#### 4.2.2 Accuracy to Estimate Spread Dynamics

Compared to the overall rate, the ability to estimate spread dynamics (see Appendix S3) decreased for all methods (Table 5). All methods, with the exception of NIR, AER, and MSD, had high rates of performance for S1 and R1 and lower rates of performance for S3 and for R3. The MSD method had a low performance rate for all scenarios and regions. CDR and MDR only had good estimation for S1 (Table 5). Similarly, CEB, AER, IAR, and COB were the best four methods for all scenarios and regions based on both R^2^ and RMSE (Table 5), whereas CEB showed consistently accurate estimates for all regions and scenarios (R^2^> 0.75).

**Table 5.** R^2^ of estimated spread dynamics by each scenario and region, and R^2^ and root mean square error (RMSE) of estimated spread dynamics for all regions and scenarios

| Spread |  | Regression Method |  |  |  |  | Boundary Displacement |  | MSD |
| --- | --- | --- | --- | --- | --- | --- | --- | --- | --- |
|  |  | CDR | IAR | MDR | NIR | AER | CEB | COB |  |
| $R^2$ by scenario | S1 | 0.83 | 0.84 | 0.81 | 0.36 | 0.84 | 0.90 | 0.83 | 0.24 |
|  | S2 | 0.60 | 0.79 | 0.45 | 0.35 | 0.89 | 0.80 | 0.77 | 0.22 |
|  | S3 | 0.05 | 0.57 | 0.06 | 0.24 | 0.63 | 0.77 | 0.58 | 0.07 |
| $R^2$ by region | R1 | 0.37 | 0.96 | 0.39 | 0.46 | 0.89 | 0.97 | 0.96 | 0.27 |
|  | R2 | 0.31 | 0.87 | 0.32 | 0.48 | 0.92 | 0.90 | 0.88 | 0.24 |
|  | R3 | 0.31 | 0.57 | 0.31 | 0.38 | 0.65 | 0.77 | 0.55 | 0.30 |
| All spread dynamics | $R^2$ | 0.33 | 0.76 | 0.34 | 0.24 | 0.80 | 0.86 | 0.76 | 0.19 |
|  | RMSE | 7.44 | 3.86 | 7.25 | NA | 3.50 | 3.30 | 4.13 | 8.75 |

S1, S2, and S3 represent symmetric spread, asymmetric spread, and long distance jump dispersal, respectively.

### 4.3 Impact of County Size and its Variation on Estimation of Spread Rate

Significantly positive and negative correlations existed between the county size and spread rates estimated by MSD and NIR, respectively (Table 6). Additionally, larger county sizes also led to higher estimated spread dynamics for IAR, CEB, and COB (Table 6). For the accuracy of estimated rates, significantly negative correlation of R^2^ with mean and CV of county size was only observed on the IAR method for overall rate, but were observed on IAR, CEB, and COB for spread dynamics (Table 6). These negative correlations suggest that accuracy is negatively impacted by the county size and its variation. However, for NIR and AER the variation in mean county size among all periods is more influential than the size of county on the estimation accuracy.

**Table 6.**
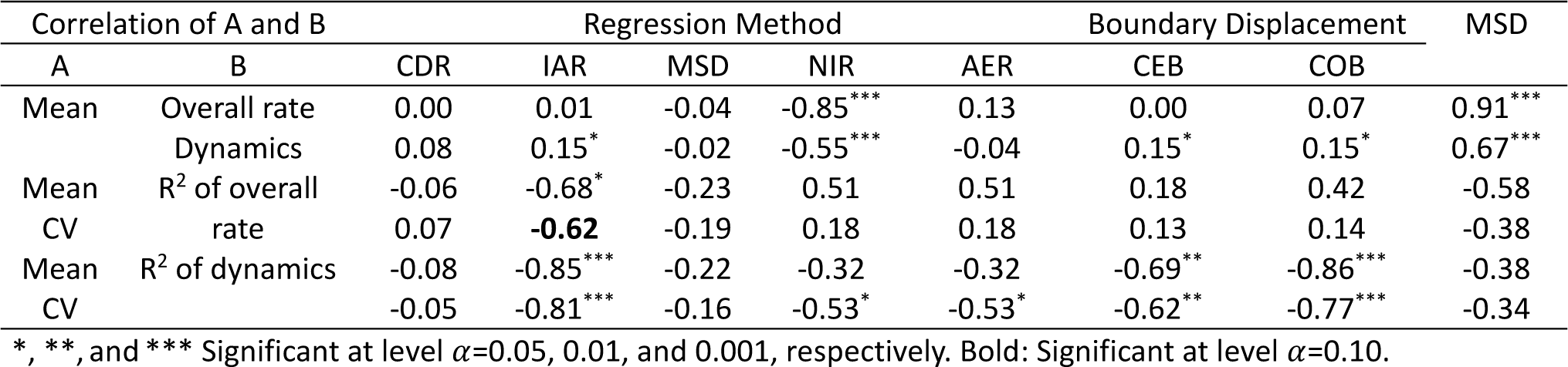
Correlation coefficient between mean of county size and estimated overall rate and spread dynamics, and between mean/coefficient of variation of county size and R^2^ for each region, spread type, and spread scenario

### 4.4 Similarity of All Methods

#### 4.4.1 Similarity of Overall Rate

All methods estimated highly correlated overall rates in R1 (r>0.90, Fig. 3 (a)) and R2 (r>=0.85, Fig. 3 (b)), whereas the large county size in R3 only dramatically decreased the similarities of MSD with the remaining methods (Fig. 3 (c)). For both S1 and S2, all methods (except NIR and MSD) estimated highly correlated overall rates among each other and with the simulation (r>=0.90, Fig. 3 (e), (f)). Similarities of CDR and MSD with other methods also decreased for S3, however very high positive correlations were still observed among IAR, AER, CEB, and COB (Fig. 3 (g)).

**Fig. 3.**
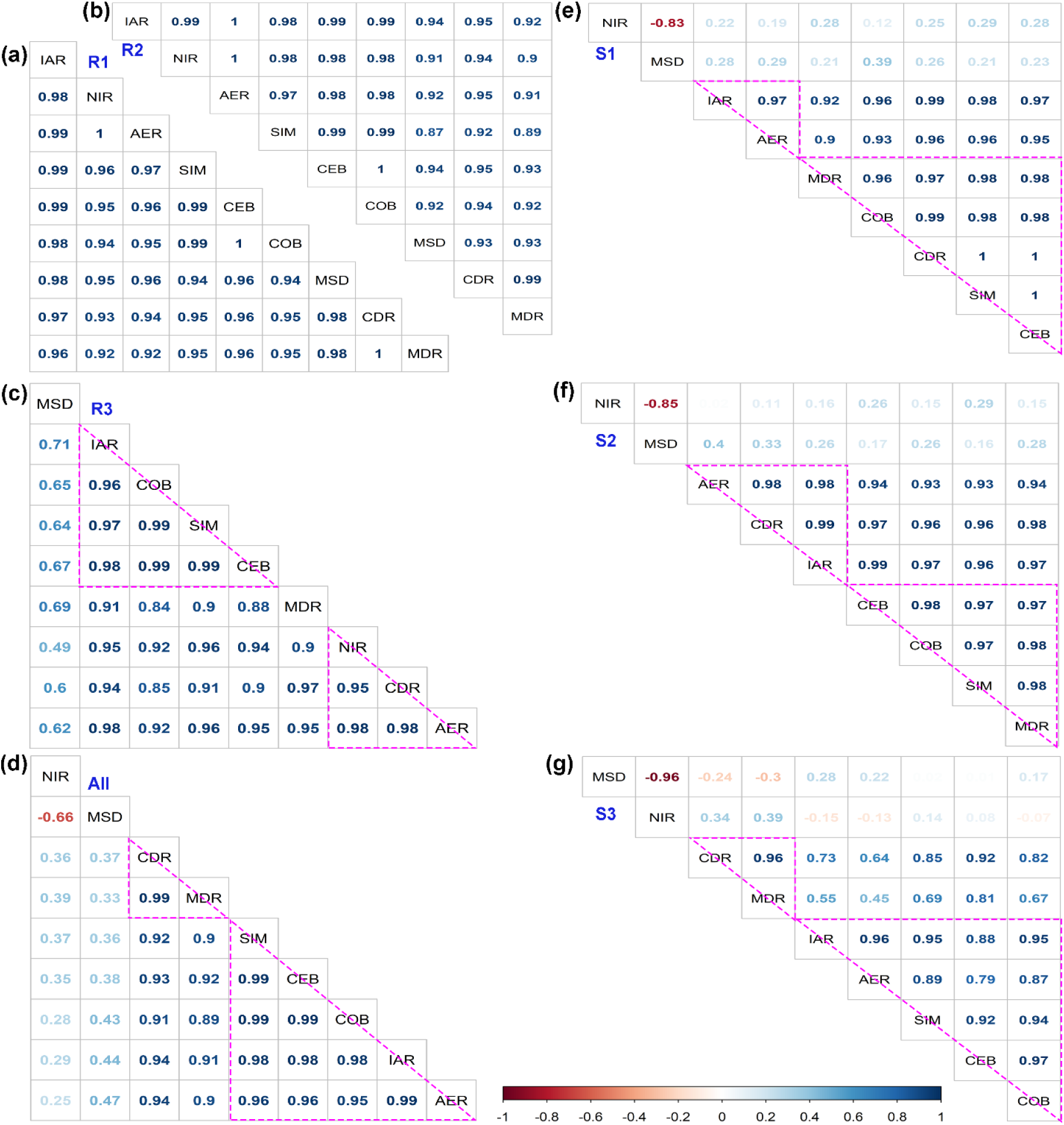
Correlation matrix (CM) and similarity pattern (SP) of estimated overall rate by all methods. **(a)-(d)** CM and SP in region 1 (R1), region 2 (R2), region 3 (R3), and all regions for three spread scenarios, respectively; **(e)-(g)** CM and SP of symmetric spread (S1), asymmetric spread (S2), and long-distance jump dispersal (S3) for all regions, respectively. Methods that are enclosed in the same triangle are classified in the same group by hierarchical clustering based on similarity of estimated rate.

#### 4.4.2 Similarity of Spread Dynamics

The spread dynamics are more sensitive to the irregularities and stochastic events than overall rate, consequently, high correlations among all methods were only observed for S1 in R1 (Fig. 4 (a)). With the increase of anisotropy and stochasticity in spread in R1 and R2, the similarities of the CDR, MDR, MSD, and NIR decreased with other methods (Fig. 4 (b)-(c)). Nevertheless, estimated dynamics by AER, IAR, COB, and CEB still had high correlations among each other and with the simulated dynamics (r>0.90, Fig. 4 (b)-(c)). The similarity of all methods was further weakened in R3, and very high correlations were only observed among IAR, COB, and CEB (r>0.80, Fig. 4 (h)).

**Fig. 4.**
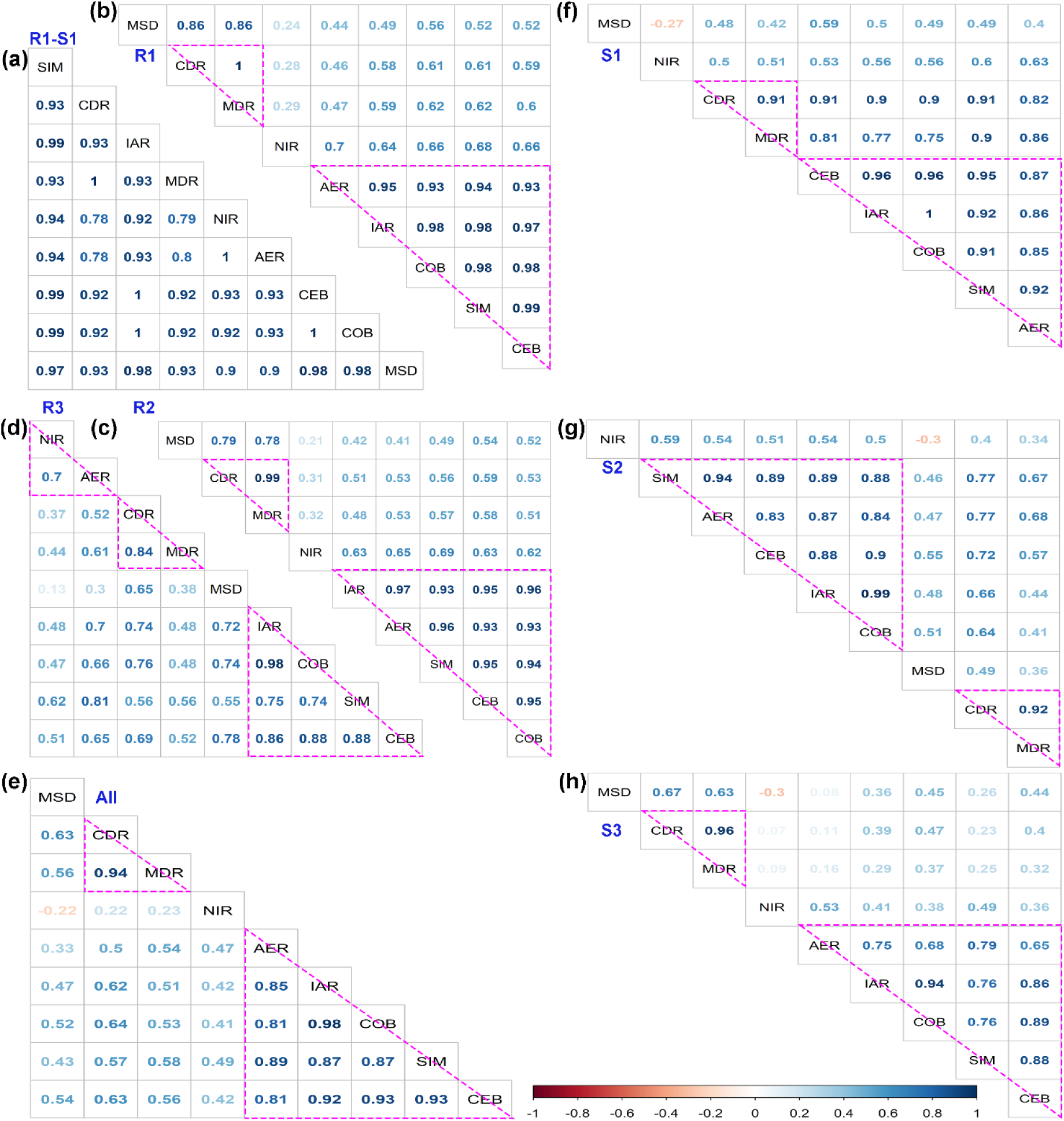
Correlation matrix (CM) and similarity pattern (SP) for estimated spread dynamics by all methods. **(a)** CM in region 1 (R1) for symmetric spread (S1)**; (b)-(e)** CM and SP in R1, region 2 (R2), region 3 (R3), and all regions for three spread scenarios, respectively; **(f)-(g)** CM and SP of S1, asymmetric spread (S2), and long distance jump dispersal (S3) for all regions, respectively. Methods that are enclosed in the same triangle are classified in the same group by hierarchical clustering based on similarity of estimated rate.

#### 4.4.3 Overall Similarity

The CDR and MDR always estimated highly correlated overall spread rate and spread dynamics (r>0.95, Fig. 3 & 6), whereas MSD and NIR always estimated low similarity with other methods when spread in all regions was analyzed. For the similarity patterns of all estimations, all methods (except NIR and MSD) estimated highly correlated overall spread rates (Fig. 3 (d)), whereas IAR, AER, CEB, and COB estimated highly correlated spread dynamics (Fig. 4 (e)). These strong correlations indicated that the estimated spread patterns were essentially similar. IAR, CEB, COB and AER were constantly classified into one group based on their similarities of estimating overall spread rate and spread dynamics (Fig. 3 & 4).

### 4.5 Case Study on Kudzu Bug

The similarity patterns of all methods for estimating spread of kudzu bug showed consistent results with the simulated study. CDR and MDR estimated highly correlated overall spread rate and spread dynamics (r>0.90, Fig. 5 (b)-(c)), meanwhile the AER, IAR, COB, and CEB were also classified into one group (Fig. 5 (b)-(c)). The spread of kudzu bug showed an asymmetric spread, thus the overall rate by MSD and spread dynamics by CDR, MDR, and MSD were less similar to the remaining methods as the same shown for S2 (Fig. 3 (f) & Fig. 4 (g)). The infested area of kudzu bug in the U.S. is similar with R1, where county size is relatively small and uniform. Consequently, the NIR method also estimated a similar spread pattern with AER, IAR, COB, and CEB as shown in R1 (r> 0.80, Fig. 3 (a), Fig. 4 (a), & Fig. 5 (c)).

**Fig. 5.**
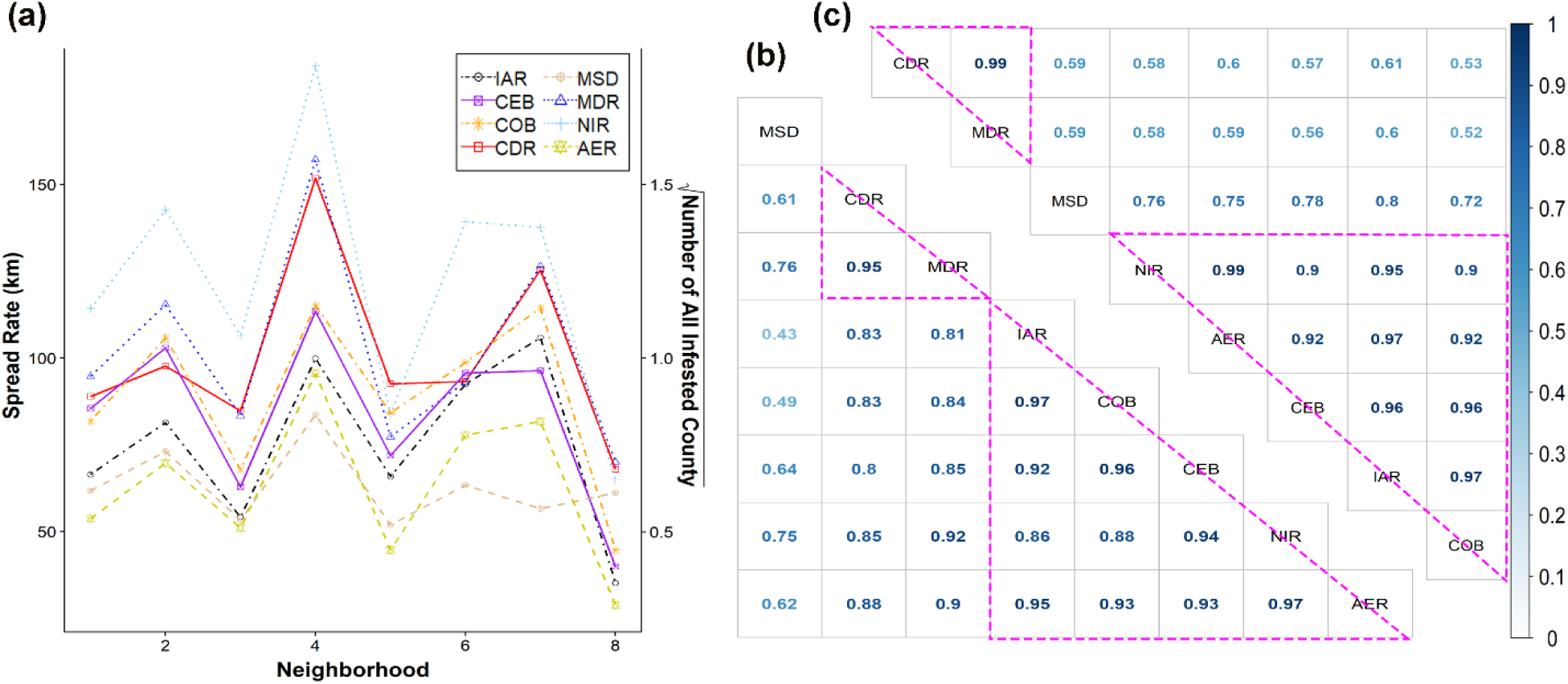
**(a)** Overall spread rate estimated by all methods in eight neighborhoods; **(b)** and **(c)** correlations of estimated overall spread rate and spread dynamics, respectively, by all methods for eight neighborhoods.

## 5 Discussion

Estimating spread is critical for prediction and early management of invasive species and understanding important factors affecting the spread (Liang et al., 2019; Paini et al., 2016; Stohlgren & Schnase, 2006). This research evaluated the ability and similarity of commonly-used methods to estimate spread with geopolitical-unit level records. Findings of this research can guide selection of optimal methods to estimate invasions with both geopolitical-unit level data and other types of invasion records. It is worth noticing that the spread rate we aimed to address in this research refers to the spread distance in a given period rather than the area expansion rate.

### 5.1 Ability of Common Methods to Estimate Spread with Geopolitical-Unit Level Records

Regression methods have long been used to determine spread patterns of invasive species (e.g., Liang et al., 2019; Liebhold et al., 1992; Mineur et al., 2010; Perrins et al., 1993). We found the cumulative value of boundary displacement methods can also be used to estimate spread patterns. The CEB, NIR, and AER always estimated the correct spread patterns regardless of variations in county size or anisotropy and stochasticity in spread.

Compared to the overall spread rate, spread dynamics can provide more complete knowledge on the spread of invasive species and better facilitate further analysis or management (Hastings et al., 2005; Liang et al., 2019). However, the estimation of spread dynamics is more challenging than the estimation of overall spread rate. Boundary displacement and MSD have been commonly used to estimate spread dynamics of invasive species (e.g., Horvitz et al., 2017; Sharov et al., 1999; Wang & Wang, 2006). Tobin et al. (2007) concluded that the boundary displacement method is better than the regression method, as the former method can describe dynamics of spread rate. In fact, two regression methods, IAR and AER, were among the top performing methods, CEB, COB, IAR, and AER, in our simulation, estimating both overall spread rate and spread dynamics accurately. For spread with clear infestation outlines, CEB can be a top choice as it constantly had the best estimation. For spread without clear infestation outlines, IAR and AER can be alternatives to estimate the spread dynamics.

Generally, the abilities of all methods to estimate invasion rates decreased with the increase of anisotropy and stochasticity in spread. However, the distance-based regression methods (i.e., CDR and MDR) were more sensitive to these irregularities and stochasticity than the area-based regression methods (i.e., IAR and AER) and boundary displacement methods. This sensitivity of distance-based regression methods is caused by the fact that their measurements can be more easily skewed by unrepresentatively long distances caused by stochasticity or heterogeneity. Consequently, CDR and MDR had low abilities to estimate overall spread rate for LDJD, and can only estimate spread dynamics well without high asymmetry.

Researchers have used MSD to estimate spread dynamics (e.g., Aikio et al., 2010; Horvitz et al., 2017) and NIR to estimate overall rates and spread patterns of invasive species (Perrins et al., 1993; Pyšek et al., 2008). However, when evaluating the accuracy across different regions, NIR and MSD constantly showed low abilities of estimating both overall rate and spread dynamics due to their sensitivity to the size of geopolitical units. Additionally, MSD constantly showed the lowest ability to estimate both overall rate and spread dynamics, and NIR had a low ability estimating spread dynamics among all methods considered. Therefore, based on this research, MSD is not recommended to estimate spread of invasive species, whereas NIR is only suggested to estimate overall spread rate when the mean sizes of geopolitical units across different periods or regions are relatively uniform. AER is a better alternative to the NIR as it rectifies the NIR by the mean area of geopolitical unit.

### 5.2 Estimating Spread with Anisotropy and Stochasticity with Geopolitical-Unit Level Records

Compared to LDJD, asymmetric spread caused by spatial heterogeneity does not seriously challenge the ability of all methods to estimate overall spread rate. Meanwhile boundary displacement methods and area-based regression methods can still have good estimation of spread dynamics under this scenario. However, when the spread is highly asymmetric and the study area is large, such as regional or larger scales, estimating rates by considering the whole infested region as one area does not reveal the spatial dynamics of spread caused by heterogeneities (Andow et al., 1990; Liang et al., 2019). To better reveal localized spread dynamics for asymmetrical spread, Andow et al. (1990) first proposed to divide the large infested area into multiple neighborhoods to increase homogeneity within each neighborhood. The neighborhood measurement had been applied in multiple research for better estimation of spread (e.g., Fraser et al., 2015; Liang et al., 2019; Morin et al., 2009). Liang et al. (2019) used a quantitative method, spatial constrained clustering, to classify a large heterogeneous region into environmentally homogeneous sub-regions. As suggested by this research, dividing a highly asymmetrical spread into several relatively symmetric spreads within each sub-region can improve the estimation accuracy. Meanwhile the neighborhood measurement also contributes to better understanding of the spread dynamics and facilitates analysis of spatial factors impacting the spreads (Liang et al., 2019). We, therefore, suggest the use of neighborhood measurement when the spread is highly asymmetric.

LDJD is often caused by rare random events and human-related activities (Nathan, 2006; Suarez et al., 2001). However, despite its rarity and stochasticity, it greatly facilitates the spread of invasive species and can be more influential than local dispersal for some species (Gilbert et al., 2004; Kot et al., 1996; Nathan, 2006; Shigesada et al., 1995; Suarez et al., 2001). For LDJD, estimation of spread rate could not indicate the actual dispersal ability of the species, but rather a rate impacted by stochastic events. LDJD causes new spread origins and obscure the spread patterns (Mineur et al., 2010; Nathan, 2006). To better estimate the ability of an invasive species to spread, researchers could set multiple spread origins, including the ones caused by LDJD, when the spread originating from LDJD are recognizable (Muirhead et al., 2006; Suarez et al., 2001). Additionally, Goldstein et al. (2019) developed a new method which could estimate the infestation boundary and detect possible LDJD with geopolitical-unit level records. However, when the spread originating from LDJD is not recognizable, boundary displacement and area-based regression methods could be preferred, as they are less sensitive to the LDJD than distance-based regression methods. Methods that estimate local spread rate determined by dispersal ability of invasive species and long-distance spread rate impacted by human-related activities can also be used. Meentemeyer et al. (2011) estimated both the short-distance and long-distance spread rate using Markov chain Monte Carlo with local and regional spread data. Suarez et al. (2001) used two different sets of data to estimate the local and regional spread rate of Argentine ants.

### 5.3 Similarity of All Methods to Estimate Spread with Geopolitical-Unit Level Record

A variety of measurements, such as spread distance and number of infested geopolitical units, are used by different methods, thus the estimated spread rates may vary based upon the measurements. However, the spread patterns, i.e., the spatial or temporal spread dynamics, revealed by different methods could be essentially similar (Liang et al., 2019; Weber, 1998). In some situations, such as determining important factors affecting spreads of invasive species (Lantschner et al., 2014; Liang et al., 2019; Sharov et al., 1999), the accuracy of estimated spread patterns matters more than the values of spread rates.

When the spread is symmetrical and the county size is relatively uniform, estimates of spread rates among all eight methods are similar. When irregularities and stochasticity occur, IAR, CEB, COB, and AER estimated similar spread dynamics, among which IAR, CEB, and COB are more similar with each other than with AER. When the overall spread rate is the question of interest, except NIR, MSD, and COB, all other five methods estimated similar overall spread rates with the simulations. Unsurprisingly, the two distance-based regression methods, CDR and MDR, always estimated highly correlated overall rates and spread dynamics. There is also no significant difference of accuracy on estimated overall rate (*P*=0.49) and spread dynamics (*P*=0.48) between these two methods. The two boundary displacement methods always estimated very highly correlated overall spread rate and spread dynamics (r>0.85), among which CEB had higher accuracy on estimating spread dynamics than COB (*P*=0.01).

## 6 Conclusions

Using simulated spread data, we found that geopolitical-unit invasion records are capable of accurately estimating spread of invasive species. Both regression and boundary displacement methods can be used to estimate the spread pattern, overall spread rate, and spread dynamics of invasive species. Selection of an optimal method depends on the question of interest, anisotropy and stochasticity in spread, and variations and sizes of geopolitical units. When the question of interest is the overall rate, among the eight methods considered,, all methods except MSD and NIR can be used for spread without LDJD. The distance-based regression methods (i.e., CDR and MDR) are more sensitive to the irregularities and stochasticity in spread than the area-based regression methods and boundary displacement methods. For LDJD, boundary displacement and area-based regression methods, i.e., IAR and AER, can be used to estimate overall rates. Estimating spread dynamics is more informative but also more challenging than estimating the overall rate. Boundary displacement methods and area-based regression methods estimated spread dynamics for all scenarios most reliably. For both overall rate and spread dynamics, boundary displacement methods have the best estimations, and CEB performs the best. However, for spread without clear infestation outlines, area-based regression methods can be good alternatives. Research using geopolitical-unit level records usually have large research scales, thus we suggest using neighborhood measurement for highly asymmetric spread to better estimate invasion dynamics.

## Acknowledgements

We would like to thank Dr. Scott Stewart at the University of Tennessee for valuable discussions on this research, and the Tennessee Soybean Promotion Board for financial support.

